# Can foot placement during gait be trained? Adaptations in stability control when ankle moments are constrained

**DOI:** 10.1101/2021.08.13.456273

**Authors:** L.A. Hoogstad, A.M. van Leeuwen, J.H. van Dieën, S.M. Bruijn

**Affiliations:** Department of Human Movement Sciences, Faculty of Behavioural and Movement Sciences, Vrije Universiteit Amsterdam, Amsterdam Movement Sciences, Amsterdam, The Netherlands; Institute of Brain and Behavior Amsterdam

**Author notes:** Shared first authorship (L.A. Hoogstad, A.M. van Leeuwen).

## Abstract

Accurate coordination of mediolateral foot placement, relative to the center of mass kinematic state, is one of the mechanisms which ensures mediolateral stability during human walking. Previously, we found that shoes constraining ankle moments decreased the degree of foot placement control with respect to the center of mass kinematic state. As such, ankle moment constraints can be seen as a perturbation of foot placement. Direct mechanical perturbations of the swing leg trajectory can improve the degree of foot placement control as an after-effect. Here, we asked whether constrained ankle moments could have a similar effect. If confirmed, this would offer a simple training tool for individuals with impaired foot placement control. Participants walked in three conditions; normal (baseline) while wearing shoes constraining ankle moments (training) and normal again (after-effects). The degree of foot placement control was calculated as the percentage of variance in foot placement that could be predicted based on the center of mass kinematic state in the preceding swing phase. During training, the degree of foot placement control decreased initially compared to baseline, but it gradually improved over time. In the after-effect condition, it was higher than during baseline, yet not significantly so. During training, we observed increased step width, decreased stride time and reduced local dynamic stability. In conclusion, constraining ankle moment control deteriorates the degree of foot placement control. A non-significant trend towards an improved degree of foot placement control after prolonged exposure to constrained ankle moments, allows for speculation on a training potential.

## 1. Introduction

Stable gait is crucial for activities of daily living. With aging, changes occur that affect the ability to maintain stability (Rubenstein, 2006; Vandervoort, 2002) and this constitutes a public health concern. Falls are common in older adults and can have severe impacts on their functionality and well-being (Rubenstein and Josephson, 2006). Furthermore, injuries related to falls lead to accumulating healthcare costs (Florence et al., 2018; Polinder et al., 2016).

Gait stability requires the coordination of the center of mass (CoM) relative to the base of support (Bruijn and van Dieën, 2018). Different strategies can be used to establish appropriate coordination between the CoM and the base of support (Reimann et al., 2018). Accurate foot placement relative to the body’s kinematic state, seems to be the dominant strategy (Bruijn and van Dieën, 2018). Foot placement defines the borders of the base of support, and through foot placement the base of support can be enlarged or displaced to accommodate the ongoing movement of the CoM (Hof, 2007; Reimann et al., 2018).

Previous work showed that foot placement can be predicted by the full body CoM kinematic state during the preceding swing phase (Hurt et al., 2010; Mahaki et al., 2019; van Leeuwen et al., 2020; Wang and Srinivasan, 2014). The degree of foot placement control can thus be quantified as the predictability of foot placement based on CoM kinematic state.

This degree of foot placement control appears to be important to maintain stability. Following a stroke, individuals demonstrate a lesser degree of foot placement control and a less stable gait (Dean and Kautz, 2015). Similarly, older adults show a lesser degree of foot placement than young adults (Arvin et al., 2018; Hurt et al., 2010), which may contribute to their elevated fall risk. Therefore, training interventions targeting foot placement control may improve gait stability. Recent research showed that in young adults, mechanical perturbations of foot placement caused an improved degree of foot placement control as an after-effect (Heitkamp et al., 2019; Reimold et al., 2020).

In previous work, we found that walking with constrained ankle moments, wearing a customized shoe (LesSchuh), decreased the degree of foot placement control (van Leeuwen et al., 2020). The LesSchuh is a shoe on which it is almost impossible to use mediolateral ankle moment control, due to a narrow ridge attached to the shoe’s sole. The decreased degree of foot placement control with constrained ankle moments may be caused by an inability to use stance leg ankle moments to control the subsequent foot placement. During targeted stepping, stance leg ankle moments contribute to accurate foot placement (Zhang et al., 2020). Prior to adjusting the swing foot to be placed on a more lateral/medial target, the center of pressure (CoP) is adjusted in the opposite direction (Zhang et al., 2020). This CoP shift seems to assist in accelerating the CoM and swing leg towards the foot placement target. Possibly, ankle moment control is also executed during steady-state walking, to facilitate control over mediolateral foot placement. Indeed, during steady-state walking an inverse relationship exists between ankle moment control and subsequent foot placement (Fettrow et al., 2019), which could reflect a similar mechanism as in targeted stepping (Zhang et al., 2020). Moreover, proper control over the swing leg can only be achieved if there is a stable base of support (i.e. the stance leg). If the stance leg is unstable, this instability can disrupt foot placement control. When wearing LesSchuh, the narrow base of support may induce such instability. Therefore, constrained stance leg ankle moments can be seen as perturbations of foot placement of the contralateral leg. Considering the evidence for after-effects of direct mechanical perturbations of the swing leg to alter foot placement (Reimold et al., 2020) prolonged exposure to ankle moment constraints may lead to an increased degree of foot placement control in walking.

The purpose of the present study was to investigate whether the degree of foot placement control improves during and after walking with ankle moment constraints. To this end, participants walked for one training session on the LesSchuh.

We hypothesized that the ankle moment constraint imposed by the LesSchuh would cause an immediate negative effect on the degree of foot placement control (H1), followed by an increase in the degree of foot placement control throughout the training condition (H2). Furthermore, we hypothesized a positive after-effect when participants return to walking on their normal shoes (H3). Finally, we hypothesized that with time, the after-effect would decrease (H4). In addition to changes in the degree of foot placement control, we explored modulations in step width and stride frequency, as compensatory stabilizing mechanisms (van Leeuwen et al., 2020). In addition, to assess whether gait changes due to LesSchuh training affected gait stability, we calculated the local divergence exponent (Bruijn et al., 2013).

## 2. Methods

### 2.1. Participants

Healthy participants between 18 and 65 years were included. Participants with a self-reported history of (neurological) disorders that could affect the ability to walk independently were excluded. Participants signed written informed consent. The experimental design and procedures were approved by the Ethical committee (VCWE-2019-108).

### 2.2. Experimental setup

Two different shoes were used for each participant. The first shoe was a normal shoe imposing no constraints on gait. The second was a shoe which constrained ankle moments; the LesSchuh. A flexible ridge along the sole, as a limited base of support, constrained ankle moment control in the frontal plane (Figure 1). The flexible ridge allowed normal roll-off and push-off, but constrained mediolateral CoP shifts.

**Figure 1.**
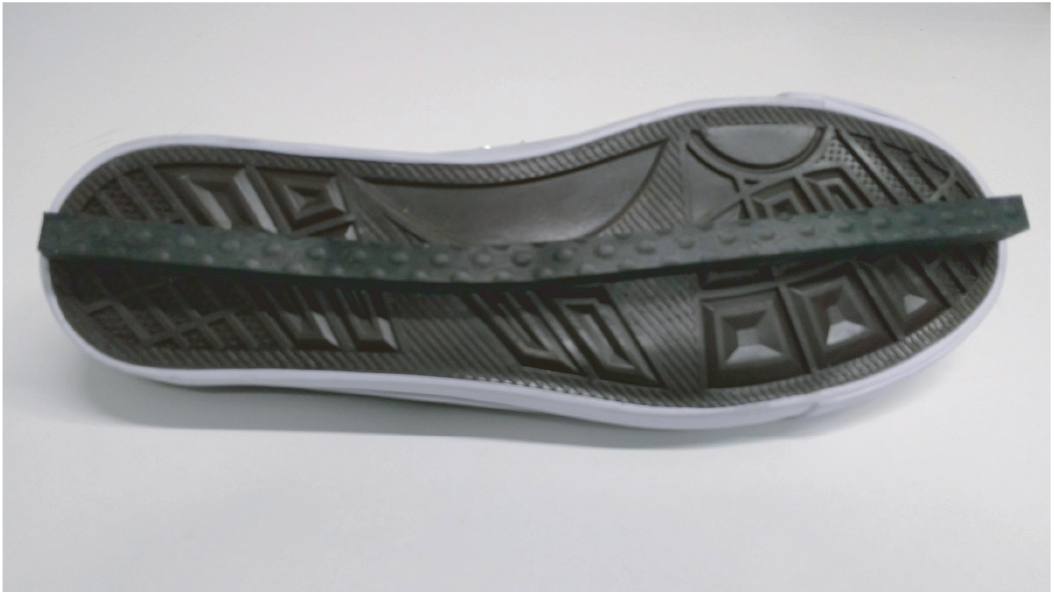
LesSchuh.

Participants walked on a split-belt treadmill with a safety bar on either side. Full body kinematics were recorded with Optotrak (Northern Digital Inc, Waterloo Ont, Canada) at a sampling rate of 50 Hz. Cluster markers of three LEDs where placed on the feet, shank, thighs, pelvis, trunk, upper arms, and forearms. 36 anatomical landmarks were digitized by using a six-marker probe.

### 2.3. Experimental design

After placing the markers, the anatomical landmarks were digitized. Subsequently, participants were asked to walk on a split-belt treadmill in three conditions (Table 1). During all three conditions the participant walked at constant (1.25 * sqrt (leg length) m/s) walking speed. In the Baseline condition, the participant walked for 10 minutes with normal shoes. In preparation of the 15-minute Training condition, the participants put on LesSchuh and the anatomical landmarks of both feet were digitized again. Then, participants were instructed to walk on the ridge of the shoe while trying to not touch the ground with the sides of the shoe’s sole. In addition, participants were instructed to keep pointing their feet straight ahead, to avoid a toeing-out strategy (Rebula et al., 2017). After the Training condition, the participant changed back to normal shoes and the anatomical landmarks were digitized again. In the After-effect condition, the participant walked for 10 minutes with normal shoes.

**Table 1.** Overview of the performed conditions

| Condition | Time | Type of shoe |
| --- | --- | --- |
| Baseline | 10 minutes | Normal shoes |
| Training | 15 minutes | LesSchuh |
| After-effect | 10 minutes | Normal shoes |

### 2.4. Data processing

Data were processed in Matlab 2021A (The MathWorks Inc., Natick, MA). Missing marker coordinates were interpolated using spline interpolation. Heel strike and toe-off events were detected from the “butterfly pattern” of the CoP derived from the force plate data (Roerdink et al., 2008). A step was defined as the period between toe-off and heel strike. Mid-swing was defined at 50 percent of the step. Step width (i.e. mediolateral foot placement) was calculated based on the position of the digitized heel markers at midstance. For estimation of the CoM, the body was segmented in 16 segments: pelvis, abdomen, thorax, head, thighs, feet, upper arms, forearms and hands. For each segment, an estimation of the mass was made based on the segment circumference and length by using a regression equation (De Leva, 1996; Zatsiorsky, 1990). The total body CoM was derived from a weighted sum of the body segments’ CoMs.

We considered two epochs of 30 strides (the first 30 and last 30 strides of each condition) for statistical analysis. Of these strides, both the left and right swing phases were included. To investigate the degree of foot placement control with respect to the CoM kinematic state, linear regression was used to predict the following foot placement based on CoM position and velocity during the preceding swing phase (Mahaki et al., 2019; van Leeuwen et al., 2020; Wang and Srinivasan, 2014). The ratio between the predicted foot placement variance and the actual foot placement variance was expressed as the relative explained variance (R^2^) and was the main outcome in this study. A high relative explained variance indicates a stronger relation between CoM state and the foot placement, i.e. a higher degree of foot placement control. We used regression equation [1] (Bruijn, 2020) in which the mediolateral CoM position (CoM_pos_) and velocity (CoM_vel_) at terminal swing predict mediolateral foot placement (FP). CoM_pos_ was defined with respect to the stance foot and both predictors were demeaned. We considered the kinematic CoM state at terminal swing, i.e. at the instant when the foot is actually being placed. This is ultimately the relationship which determines how effective foot placement will be in reversing the CoM movement. FP was defined as the demeaned mediolateral distance between the two feet. The positions of the feet were determined at midstance of the stance foot, based on the digitized heel marker. β_pos_ and β_vel_ represent the regression coefficients and ε the residual of the model (i.e. the discrepancy between predicted and FP). For each epoch of 30 strides, we determined the R^2^ of this regression.

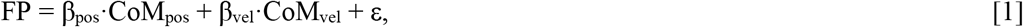

The local divergence exponent was determined (Bruijn et al., 2013; Bruijn, 2021; Mehdizadeh, 2019; Rosenstein et al., 1993) by first constructing a six-dimensional state space based on time-delayed (25 samples) copies of the 3D CoM velocity. To this end, the signal was resampled so that on average each stride was 100 samples in length. Subsequently, we tracked the divergence for each time point and its five nearest neighbors for 1000 samples. A nearest neighbor was defined as a point at the smallest Euclidian distance, at least half an average stride before or half an average stride after the current time point. We computed the average logarithmic divergence curve and fitted a line (least squares fit) to this curve over the first 50 samples. Finally, we defined the local divergence exponent as the slope of this line. The higher the local divergence exponent, the less stable the gait pattern.

Step width was defined as the mean distance between mediolateral foot placements. Stride time was defined as the time between two heel strikes of the same leg.

Lastly, the toe-out angles were checked during the baseline and training conditions. We determined the local coordinate systems of the feet based on the feet’s anatomical landmarks (calcaneus, malleoli, second toe tip). The x-, y- and z-axes represented respectively the local forward, mediolateral and vertical axes. A z-x-y Euler decomposition was performed on the corresponding orientation matrices. We considered the angle around the vertical axis at midstance as the toe-out angle. We defined the toe-out angle as zero when the feet pointed straight-ahead, positive angles to represent toeing-out, and negative angles to represent toeing-in.

The data and code for the analysis can be found online: https://surfdrive.surf.nl/files/index.php/s/ZgLVUg7ftfPkNTO, and will be uploaded to Zenodo after the paper has been accepted for publication.

### 2.5. Statistical analysis

For the statistical analyses, of each baseline, training and after-effect trial, we considered the first and last 30 strides, as the start and the end of each trial.

To check whether participants followed the instructions, we tested toe-out angles with a repeated measures ANOVA with the factor Trial (“baseline end”, “training start”, “training end”).

To test for changes during the baseline trial, we performed paired t-tests between the start and end values of the baseline trial for the R^2^, stride time, step width and local divergence exponent.

To test each of our hypotheses we performed paired t-tests on the R^2^ transformed by a modified Fisher transform. For H1, we expected a lower R^2^ at the start of the training as compared to the end of the baseline trial. In line with H2, we expected a higher R^2^ at the end of the training as compared to the start of the training. Following H3, we expected a higher R^2^ at the start of the after-effect condition as compared to the end of the baseline trial. For H4, we expected the R^2^ at the end of the after-effect condition to be lower than at the start. Moreover, we explored stride time, step width and local divergence at the same time points. For all statistical tests, *p*<0.05 was treated as significant.

## 3. Results

Nineteen healthy adults participated in this experiment. Four participants were excluded from further analysis due to measurement errors that affected the calculations. All data were normally distributed. On average participants took 529 (SD = 16) steps in the 10-minute baseline condition, 836 (SD= 35) steps in the 15-minute training condition and 525 (SD =18) steps in the 10-minute after-effect condition.

### Compliance with instructions

Participants pointed their toes more outward when walking with constrained ankle moments (training; Figure 2). The difference with baseline end was significant (p<0.05) for training end but not for training start.

**Figure 2.**
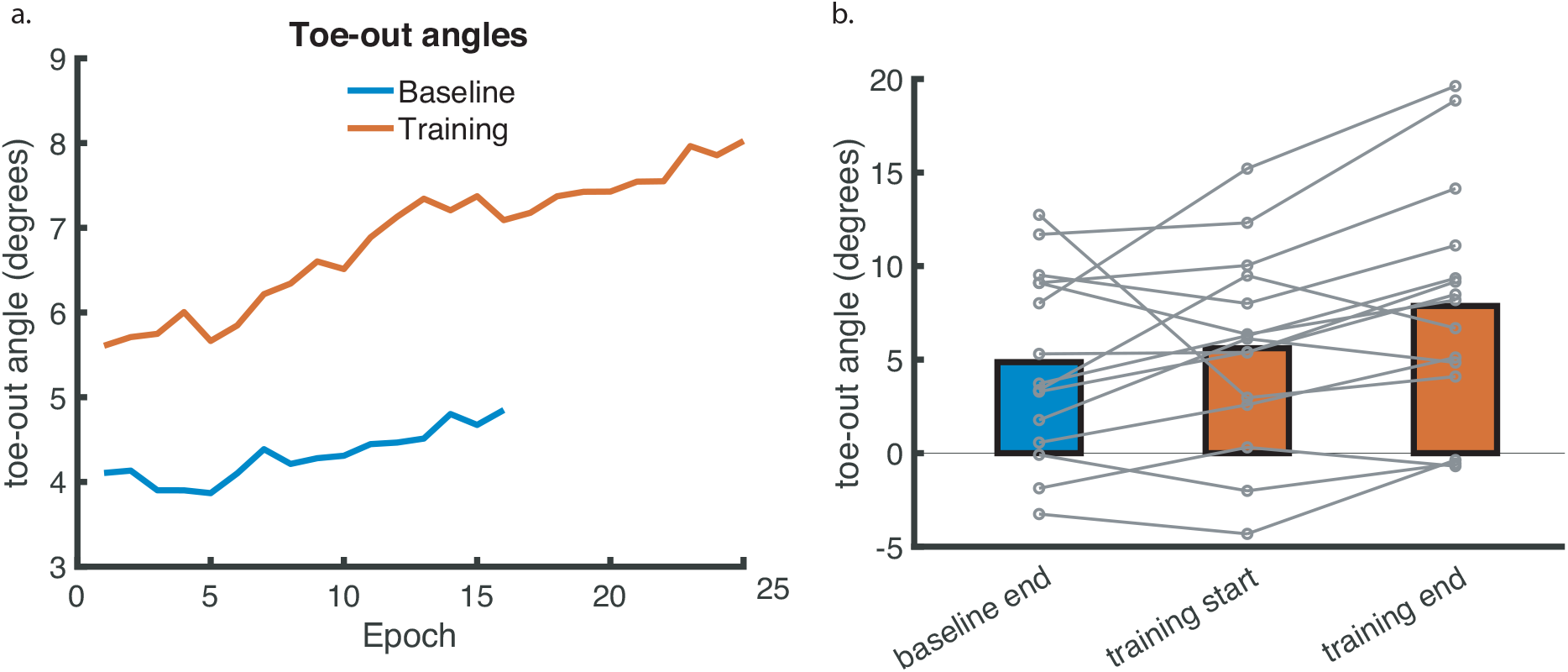
Toe-out angles. (a) Mean toe-out angles across 30 stride episodes (b) Mean toe-out angles (and individual data points in grey). The individual data points have been connected. For illustrative purposes, in figure 2a, the data is depicted into epochs of 30 steps up to the number of epochs for which all participants had a full final epoch (i.e. including 30 steps). Positive and negative angles represent respectively toeing-out and toeing-in. Zero is defined based on the foot’s local coordinate system pointing straight ahead.

### Baseline condition

Significant differences in R2, stride time and step width were found when comparing the end to the start of the baseline trial (Figure 3; *p* < 0.05). We found no significant change in gait stability over the baseline measurement (*p >* 0.05).

**Figure 3.**
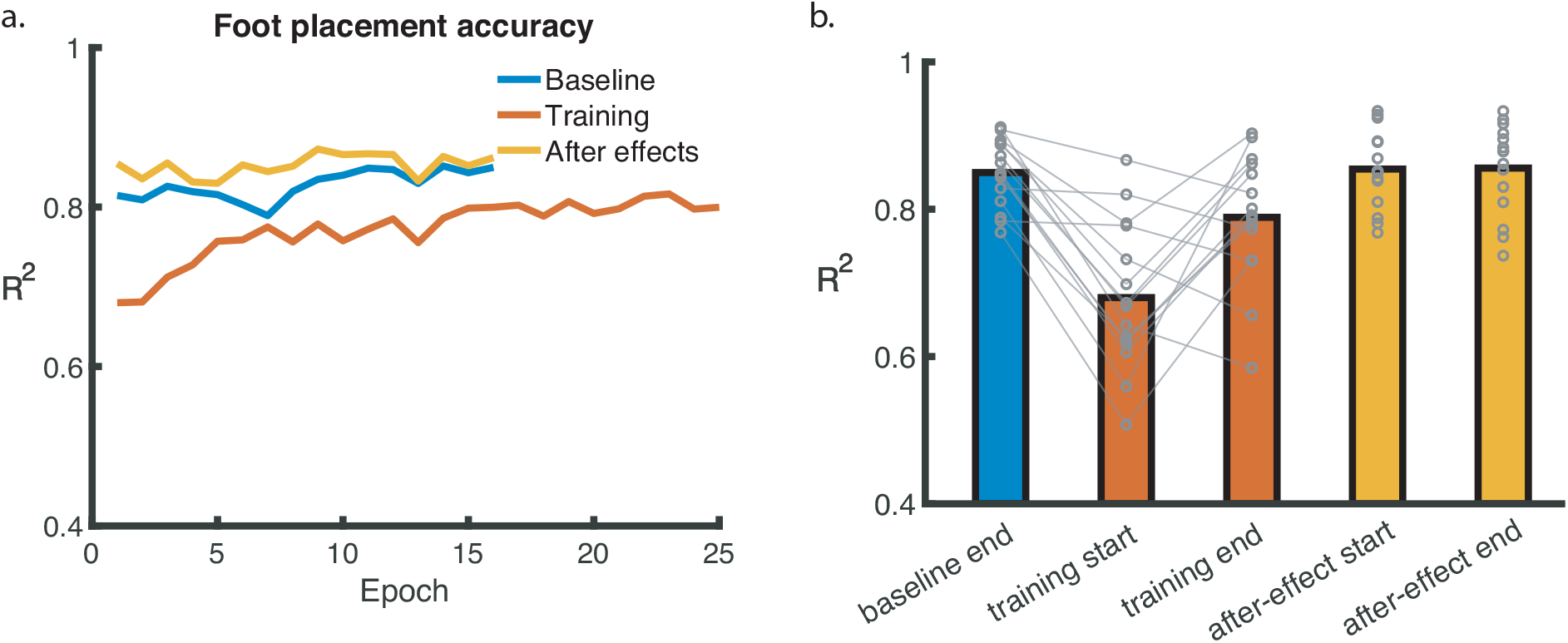
The degree of foot placement control (during the baseline, training and after-effect conditions). (a) Mean relative explained variance (R^2^) of the foot placement prediction model across 30-strides episodes, (b) Mean R^2^ (and individual data points in gray) at the end of the baseline condition, the start and end of the training condition and at the start and end of the after-effect condition. For the significant effects, the individual data points have been connected. For illustrative purposes, in figure 3a, the data is depicted into epochs of 30 steps up to the number of epochs for which all participants had a full final epoch (i.e. including 30 steps)

### Foot placement

Walking with LesSchuh led to an immediate decrease in R^2^ (*p* <0.05). A significant increase in R^2^ occurred during the training trial (Figure 3;*p* <0.05). We found no significant difference in R^2^ between the end of the baseline trial and the start of the after-effect trial (*p* > 0.05), nor a significant difference between the start and the end of the after-effect trial (*p*>0.05).

### Stride time

Stride times were significantly shorter at the start of the training trial than at the end of the baseline trials (Figure 4; *p*<0.05). During training, stride time increased, resulting in a significant difference in stride time between start and end of training (*p*<0.05). At the start of the after-effect trial, stride time was significantly shorter than at the end of the baseline trial (*p*<0.05). There was a significant increase in stride time from the start until the end of the after-effect condition (*p*<0.05).

**Figure 4.**
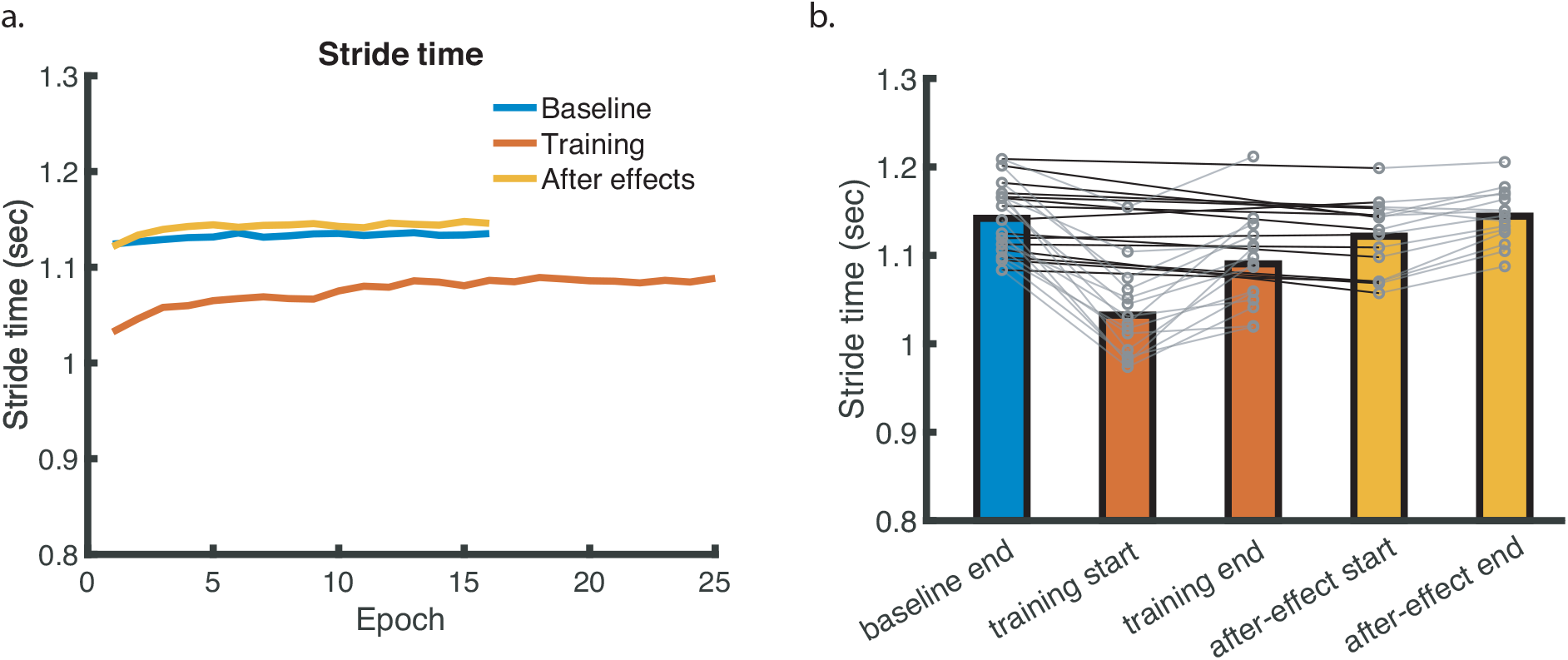
Stride times (during the baseline, training and after-effect conditions). (a) Mean stride time across stride episodes (b) Mean stride (and individual data points in gray) at the end of the baseline condition, the start and end of the training condition and at the start and end of the after-effect condition. For the significant effects, the individual data points have been connected. For illustrative purposes, in figure 4a, the data is depicted into epochs of 30 steps up to the number of epochs for which all participants had a full final epoch (i.e. including 30 steps).

### Step width

Step width was increased compared to baseline at the start of the training (Figure 5; *p*<0.05). We found no significant decrease in step width during the training (*p*>0.05), nor a significant after-effect (*p*>0.05). In line with this, there was no significant difference between the start and the end of the after-effect trial (*p*>0.05).

**Figure 5.**
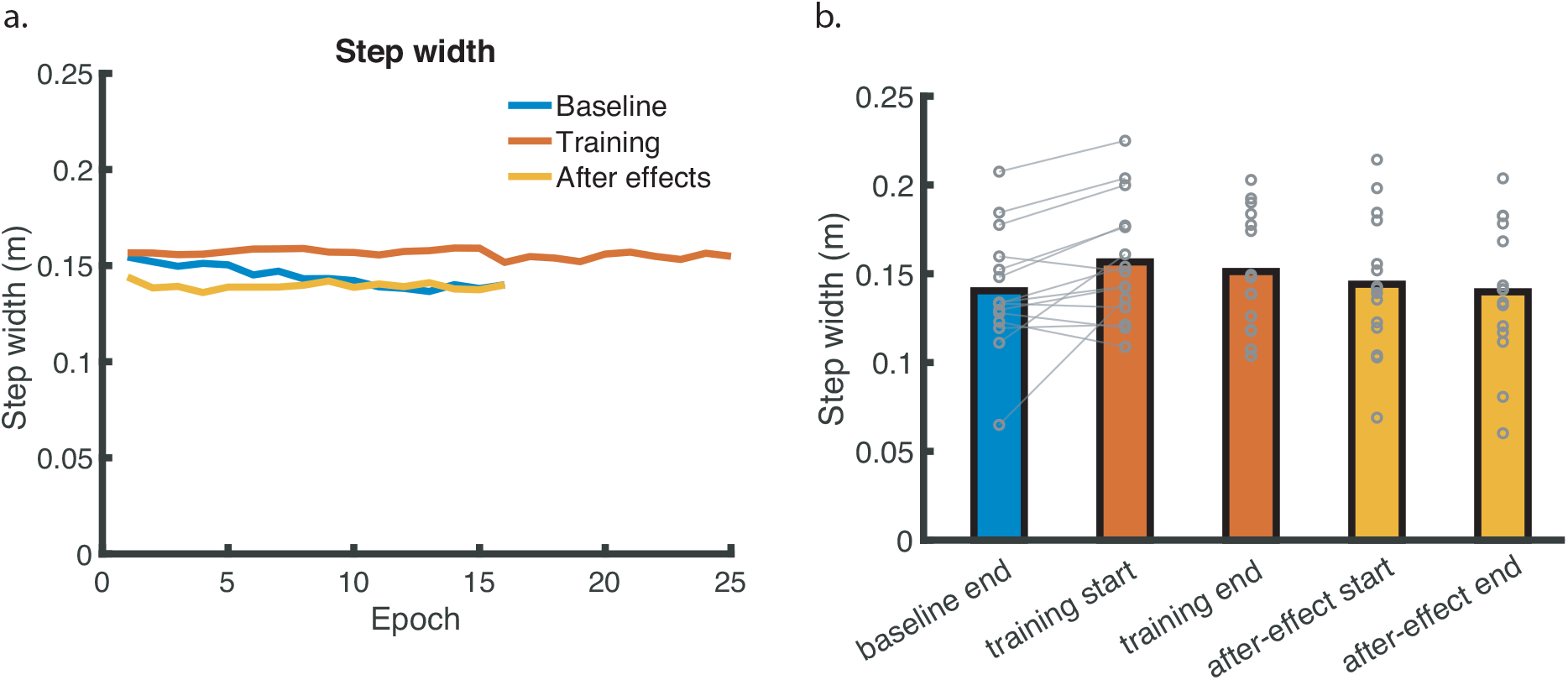
Step width (during the baseline, training and after-effect conditions). (a) Mean step width across stride episodes, (b) Mean step width (and individual data points in gray) at the end of the baseline condition, the start and end of the training condition and at the start and end of the after-effect condition. For the significant effect, the individual data points have been connected. For illustrative purposes, in figure 5a, the data is depicted into epochs of 30 steps up to the number of epochs for which all participants had a full final epoch (i.e. including 30 steps).

### Gait stability

The local divergence exponent was significantly increased (i.e. local dynamic stability decreased) at the start of training as compared to the end of the baseline trial (Figure 6; *p*<0.05). No significant change occurred during training, nor was there an after-effect (*p*>0.05). There was no change in stability between the start and end of the after-effect condition (*p>*0.05).

**Figure 6.**
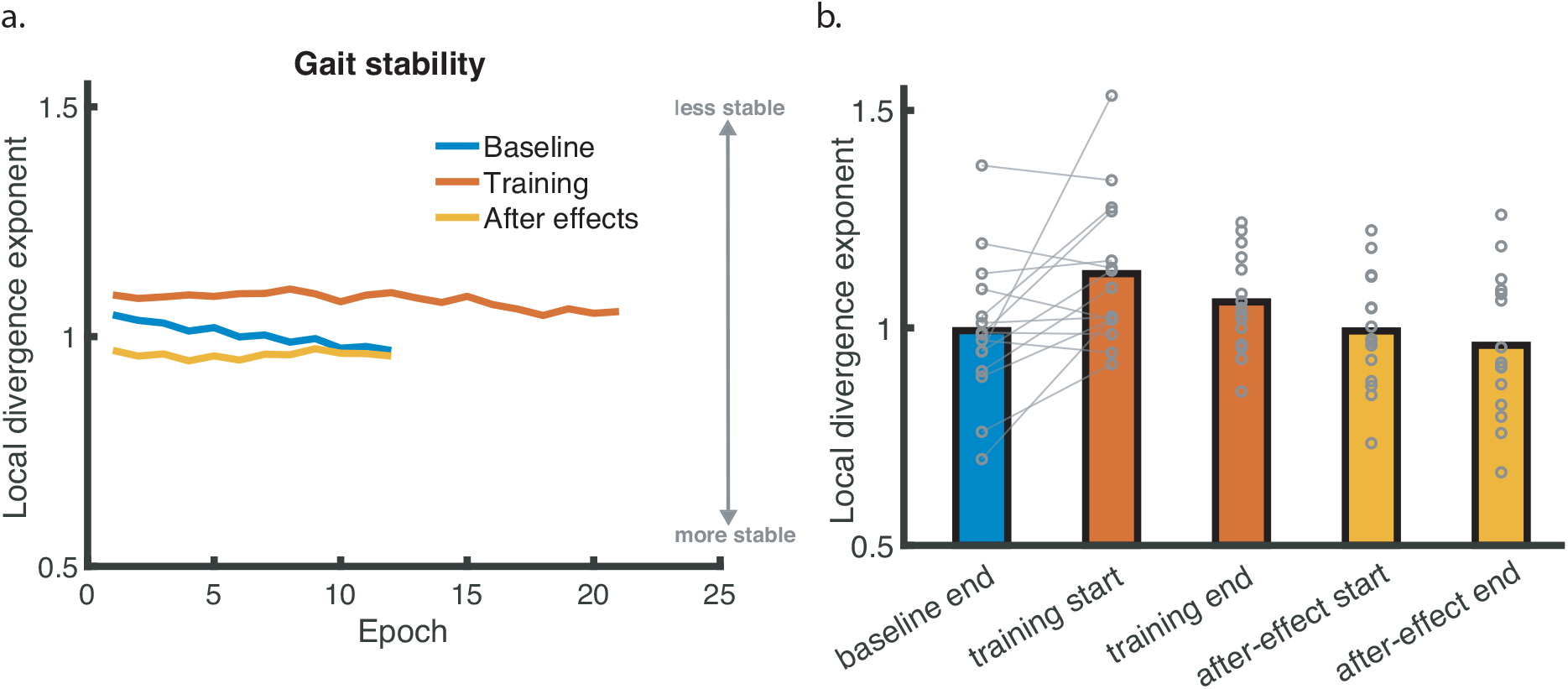
Gait stability (during the baseline, training and after-effect conditions. (a) Mean local divergence exponents across stride episodes, (b) Mean local divergence exponents (and individual data points in gray) at the end of the baseline condition, the start and end of the training condition and at the start and end of the after-effect condition. A lower local divergence exponent represents increased gait stability. For the significant effect, the individual data points have been connected. For illustrative purposes, in figure 6a, the data is depicted into epochs of 30 steps up to the number of epochs for which all participants had a full final epoch (i.e. including 30 steps).

## 4. Discussion

Our main purpose was to investigate whether the degree of foot placement control improves after walking with shoes constraining ankle moments. We used ankle moment constraints as a foot placement perturbation to train the degree of foot placement control. We verified that the degree of foot placement control decreased while walking with shoes constraining ankle moments (H1). We also observed an increasing degree of foot placement control during longer exposure to the ankle moment constraining shoes (H2). Despite this adaptation, the degree of foot placement control did not significantly increase as an after-effect upon returning to normal shoes (opposing H3). As an exploratory analysis, we assessed changes in stride time, step width and local dynamic stability. Since the degree of foot placement control is a relative measure, dependent on the absolute variance, we have added the absolute explained variance, the foot placement error and the variability in foot placement and CoM kinematic state as supplementary material. Below we discuss the gait changes per condition.

### Gait changes during the baseline condition

The degree of foot placement control, stride time and step width changed significantly from the start until the end of baseline. Such changes reflect the need for habituation to treadmill walking. 6-7 minutes (425 strides) of treadmill walking is sufficient to achieve stable gait parameters (Meyer et al., 2019). Since participants walked for 10 minutes in the baseline condition, one may reasonably assume the last 30 strides of these trials to provide a valid reference.

### Gait changes in the training condition

In the training condition, participants walked while wearing LesSchuh (Figure 1). We instructed participants to keep their feet pointing straight ahead. In this way, we intended to avoid a CoP shift due to a compensatory toeing-out strategy (Rebula et al., 2017). It seems participants tried to follow the instructions at the start of the training, but failed to do so nearing the end of the training trial (figure 2). With about three degrees toeing-out as compared to at the end of the baseline condition, this would allow for an approximate 1.3 cm CoP shift, to compensate for the ankle moment constraint (assuming a shoe length of 25 cm, we get *sin(3deg) * 25 = 1*.*3 cm*, supplementary material V). Despite this, the ankle moment constraints led to other gait changes as well. As expected from earlier results (van Leeuwen et al., 2020), the degree of foot placement control decreased. This decrease in the degree of foot placement control could be attributed to both a lesser contribution of foot placement control to mediolateral stability (lower absolute explained variance, supplementary material I) and a lower precision in foot placement control (larger foot placement error, supplementary material II). Also in line with earlier work (van Leeuwen et al., 2020), step width and stride frequency increased. Increasing step width and stride frequency appears to be an appropriate strategy to maintain stability, in spite of a reduced degree of foot placement control (Hak et al., 2013; Perry and Srinivasan, 2017). However, here local dynamic stability diminished when ankle moments were constrained, despite these compensatory mechanisms. Moreover, during unconstrained steady-state walking, step width and stride frequency seem to be largely determined by an energetic optimum (Delextrat et al., 2011; Donelan et al., 2001). Thus, increasing step width and frequency likely increases energy cost. As we have speculated earlier (van Leeuwen et al., 2020), adopting a careful gait strategy (Wu et al., 2015) could be a temporary compensatory strategy that would disappear when a certain degree of foot placement control is regained, and a more efficient step width (Donelan et al., 2001; Perry and Srinivasan, 2017) and stride time can be adopted.

Indeed, throughout the training condition, the degree of foot placement control gradually improved towards the baseline level, suggesting a training potential. This improvement reflected mainly a larger contribution of foot placement control to mediolateral gait stability (higher absolute explained variance, supplementary material I), whilst the precision in foot placement control did not improve (high foot placement error, supplementary material II). A higher degree of foot placement control is expected to allow a narrower step width while remaining equally stable (Perry and Srinivasan, 2017). However, although stride time increased again throughout the training, step width remained enlarged. Perhaps this is related to the fact that, at the end of the training condition, the degree of foot placement control, was still lower than during baseline. Possibly, the degree of foot placement control will continue to improve with longer exposure to ankle moment constrained walking, before allowing a stable, energetically efficient gait pattern.

Alternatively, it may be impossible to increase the degree of foot placement control with LesSchuh on a split-belt treadmill beyond levels found here. Recent unpublished findings on walking with LesSchuh on both a split-belt and a single-belt treadmill showed that the initial reduction in the degree of foot placement cannot be attributed to LesSchuh in itself. The reduction was the result of the combination of LesSchuh and the split-belt treadmill. On the single-belt treadmill, the degree of foot placement control did not decrease (tested for CoM kinematic state at terminal swing). As participants walked with wider steps on the split-belt treadmill, it seems that the gap between the belts imposes an additional (perceived) constraint. Apart from coordinating foot placement with respect to the CoM, the split-belt imposes the constraint of not stepping in the gap between the two belts. If similar compliance with the split-belt constraint persists over time, it may well be that, when walking with LesSchuh on a split-belt treadmill, the degree of foot placement control will never regain or exceed its baseline level. Since we have already seen degrees of foot placement control similar to baseline when walking with LesSchuh on a single-belt treadmill, the results of the current study cannot be generalized to a single-belt treadmill.

### Gait changes in the after-effect condition

Although the degree of foot placement control appeared higher throughout the after-effect trial compared to baseline (figure 3), this difference was not significant. Therefore, we cannot conclude that constraining ankle moments can lead to a training effect on the degree of foot placement control during normal steady-state walking. Still, we argue that these results suggest a training potential of walking with constrained ankle moments.

Every epoch, the average degree of foot placement control was higher in the after-effect condition as compared to the baseline condition. The same holds for the contribution of foot placement control to mediolateral stability (absolute explained variance, supplementary material I). Taken together with the increase in the degree of foot placement control during training, we speculate that walking on LesSchuh for a longer duration, may induce a (significant) larger after-effect. Alternatively, a ceiling effect may have resulted in a statistically undiscernible after-effect. Young adults already have a relatively high degree of foot placement control (Arvin et al., 2018; Hurt et al., 2010; Wang and Srinivasan, 2014), which may limit their capacity for improvement. A larger training effect could be expected in older adults (Arvin et al., 2018; Hurt et al., 2010) and patients with an impaired gait pattern (Dean and Kautz, 2015). Lastly, we speculate that training with additional constraints on other compensatory strategies, i.e. on average step width and stride frequency, may be more effective in enhancing the degree of foot placement control. If these compensatory strategies can no longer be relied upon, participants may be forced to improve their degree of foot placement control to prevent instability. Then, longer exposure to LesSchuh may not only increase the contribution of foot placement control to mediolateral stability, but perhaps also its precision. Future studies should include a control experiment during which participants walk for the same duration, but while wearing normal shoes during all trials. This could clarify whether any “after-effect” truly reflects a training potential.

## Conclusion

Walking with ankle moment constraints perturbed foot placement control. This was reflected by a decrease in the degree of foot placement control in relation to the CoM kinematic state. With longer exposure to the ankle moment constraint, participants adapted, showing an increased degree of foot placement control over time. When walking with normal shoes again, no significant after-effects were found. Nevertheless, there were indications of a training potential of walking with ankle moment constraints. It seems the interdependency of ankle moment and foot placement control provides an opportunity for training interventions.

## Acknowledgements

The authors are thankful for the participants and (technical) assistance during the experiment. We are especially grateful for Leon Schutte, who developed the LesSchuh. The Dutch Research Council (NWO) (https://www.nwo.nl/en/) funded Sjoerd Bruijn (016.Vidi.178.014) and Moira van Leeuwen.

