## Supplementary material for "Can foot placement during gait be trained? Adaptations in stability control when ankle moments are constrained"

#### I Absolute explained variance

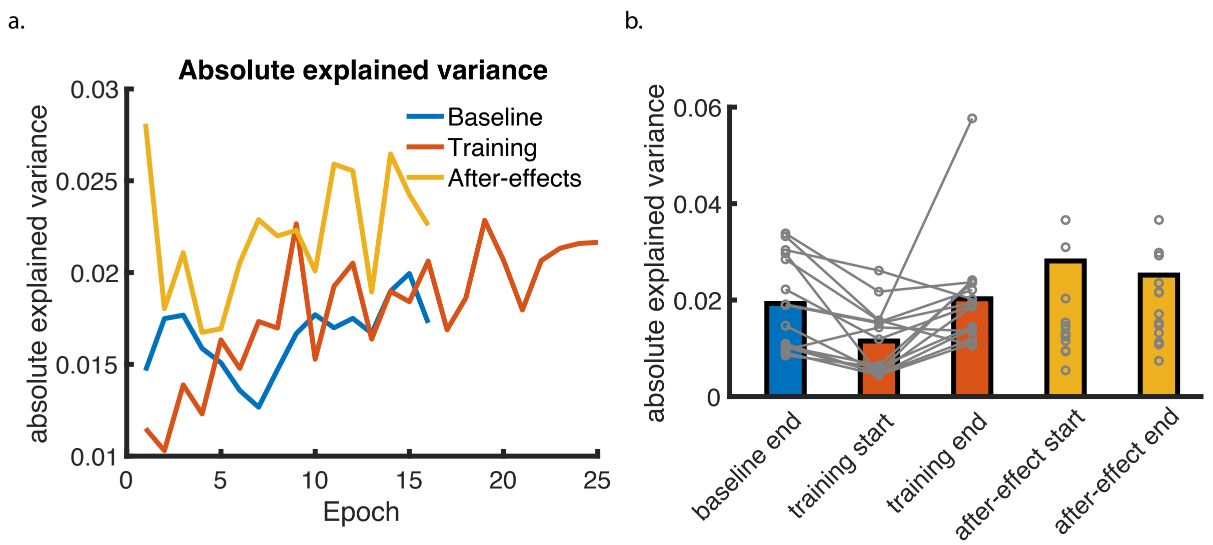


SI. Fig 1. Absolute explained variance of the foot placement model. (a) Mean absolute explained variance across 30 stride episodes. (b) Mean absolute explained variance (and individual data points in grey) at the end of the baseline condition, the start and end of the training condition and at the start and end of the after-effect condition. For significant effects of the degree of foot placement control ($R^2$, Figure 3), the individual data points have been connected. For illustrative purposes, in panel a, the data is depicted into epochs of 30 steps up to the number of epochs for which all participants had a full final epoch (i.e. including 30 steps). In accordance with the decrease in the degree of foot placement control (Figure 3) at training start as compared to baseline end, based on visual inspection, the absolute explained variance decreased. Similarly, the increase in the degree of foot placement control ($R^2$) throughout the training (Figure 3), is also reflected in an increase in the absolute explained variance.

### II Standard deviation of foot placement error

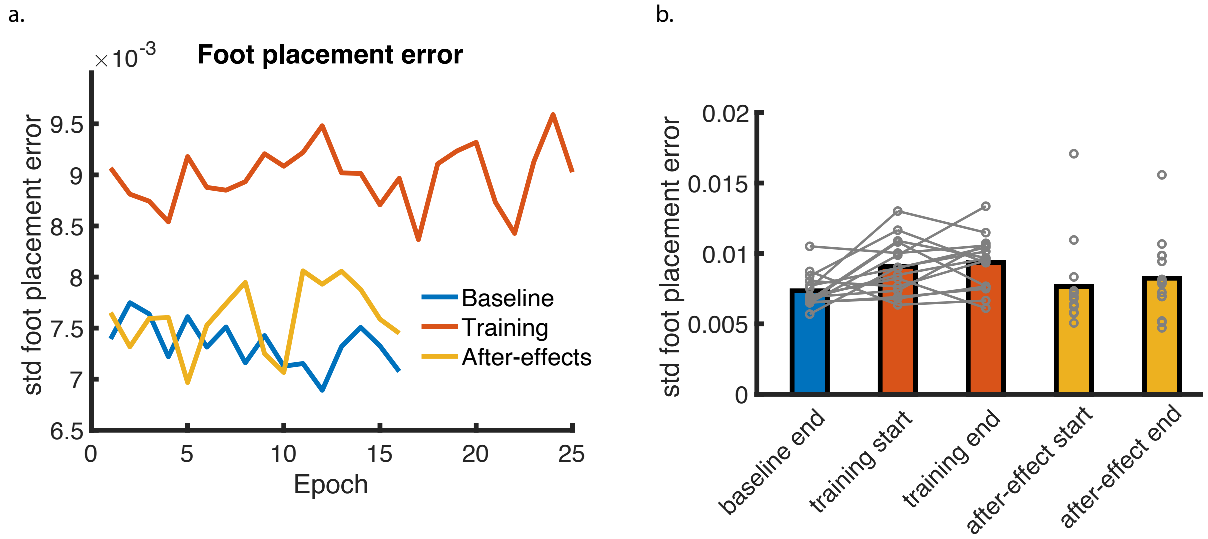


SII. Fig 1. Standard deviation of the foot placement error. (a) Mean unexplained variance across 30 stride episodes. (b) Mean unexplained variance (and individual data points in grey) at the end of the baseline condition, the start and end of the training condition and at the start and end of the after-effect condition. For significant effects of the degree of foot placement control ($R^2$, Figure 3), the individual data points have been connected. For illustrative purposes, in panel a, the data is depicted into epochs of 30 steps up to the number of epochs for which all participants had a full final epoch (i.e. including 30 steps). In accordance with the decrease in the degree of foot placement control (Figure 3) at training start as compared to baseline end, based on visual inspection, foot placement error increased. However, whereas the degree of foot placement control improves from training start to training end (figure 3), foot placement error does not decrease throughout the training trial.

### III Center-of-mass swing phase variability

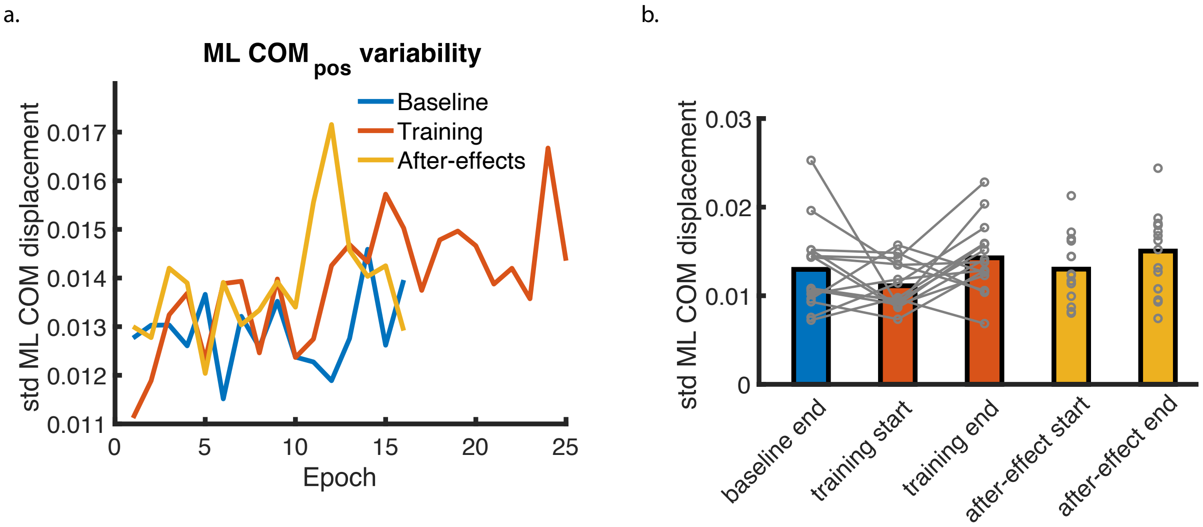


SIII. Fig 1. Mediolateral center-of-mass position ($CoM_{pos}$) variability. (a) Mean $CoM_{pos}$ variability across 30 stride episodes. (b) Mean $CoM_{pos}$ variability (and individual data points in grey) at the end of the baseline condition, the start and end of the training condition and at the start and end of the after-effect condition. For significant effects of the degree of foot placement control ($R^2$ , Figure 3), the individual data points have been connected. For illustrative purposes, in panel a, the data is depicted into epochs of 30 steps up to the number of epochs for which all participants had a full final epoch (i.e. including 30 steps). Like the degree of foot placement control (Figure 3), based on visual inspection, $CoM_{pos}$ variabilitydecreased at training start, and increased throughout the training.

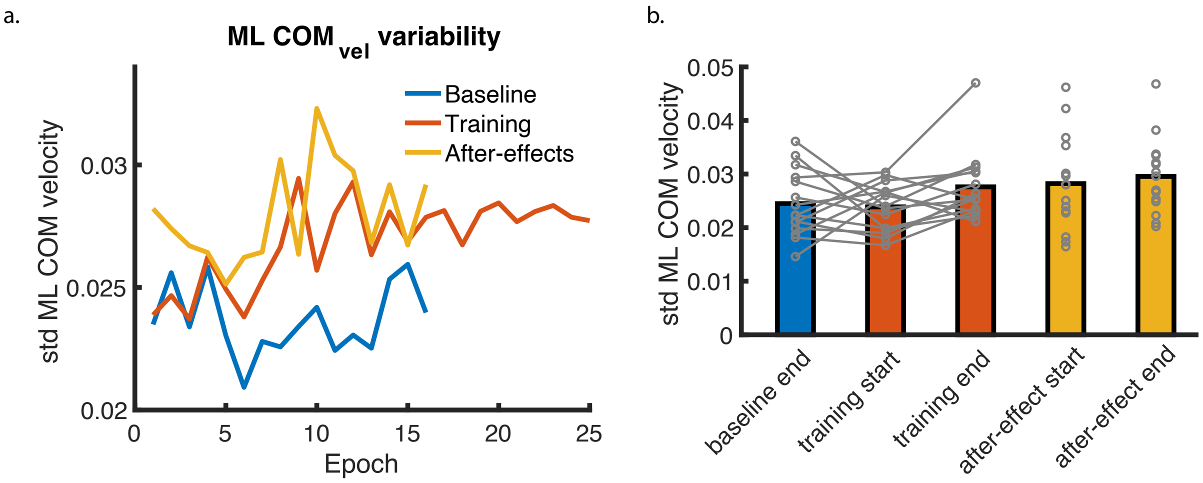


SIII. Fig 2. Mediolateral center-of-mass velocity ($CoM_{vel}$) variability. (a) Mean $CoM_{vel}$ variability across 30 stride episodes. (b) Mean $CoM_{vel}$ variability (and individual data points in grey) at the end of the baseline condition, the start and end of the training condition and at the start and end of the after-effect condition. For significant effects of the degree of foot placement control ($R^2$, Figure 3), the individual data points have been connected. For illustrative purposes, in panel a, the data is depicted into epochs of 30 steps up to the number of epochs for which all participants had a full final epoch (i.e. including 30 steps). Like the degree of foot placement control (Figure 3), based on visual inspection, $CoM_{vel}$ variability decreased at training start, and increased throughout the training.

#### IV Foot placement variability

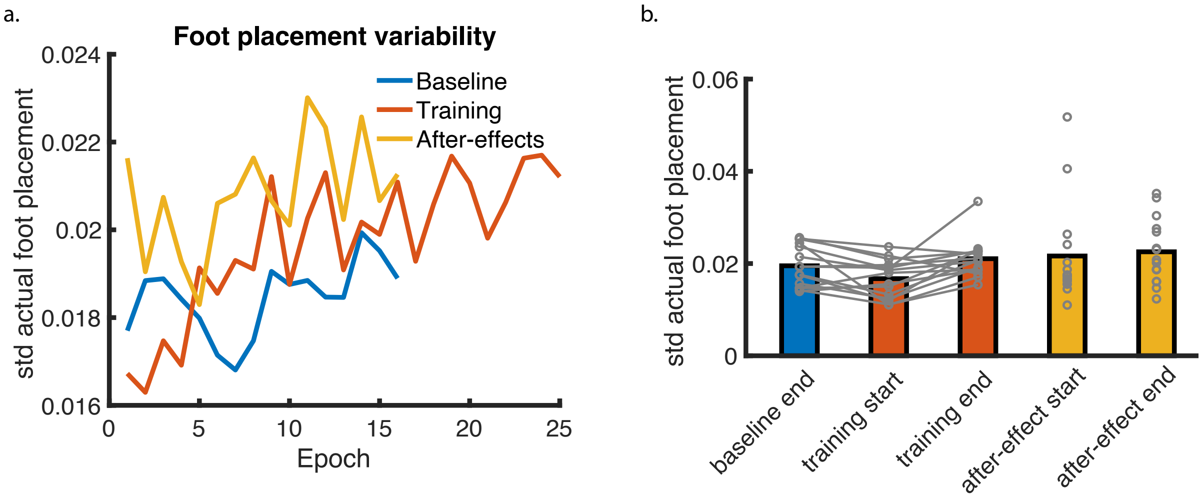


SIV. Fig 1. Foot placement variability. (a) Mean foot placement variability across 30 stride episodes. (b) Mean foot placement variability (and individual data points in grey) at the end of the baseline condition, the start and end of the training condition and at the start and end of the after-effect condition. For significant effects of the degree of foot placement control ($R^2$ , Figure 3), the individual data points have been connected. For illustrative purposes, in panel a, the data is depicted into epochs of 30 steps up to the number of epochs for which all participants had a full final epoch (i.e. including 30 steps). Like the degree of foot placement control (Figure 3), based on visual inspection, the foot placement variability decreased at training start, and increased throughout the training.

#### V Toeing-out strategy when wearing LesSchuh

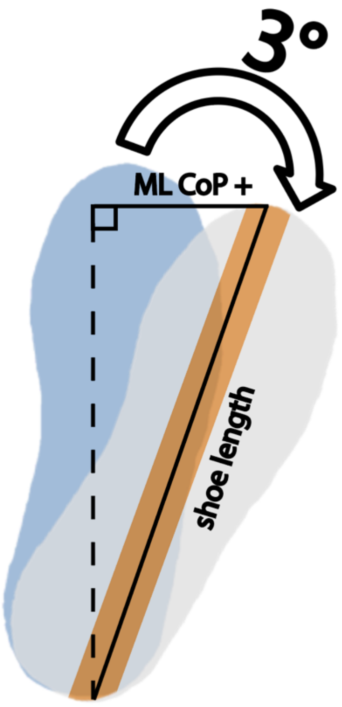


$$\text{ML CoP +} \;=\; \sin(3\ \text{degrees}) * \text{shoe length}$$
$$\text{ML CoP +} \;=\; \sin(3\ \text{degrees}) * \ 0.25 = 1.3\ \text{cm}$$

SV. Fig1. Estimated additional mediolateral center of pressure shift (ML CoP+) due to toeing-out. Despite the instructions, at the end of the training condition, on average participants walked with more toeing-out, as compared to at baseline end. There is about three degrees between the mean toe-out angle at training end, as compared to the mean toe-out angle at baseline end. In this figure we show that with an estimated shoe length of 25 centimeters, these three degrees would lead to an approximate 1.5 additional centimeters that the center of pressure can shift laterally, whilst remaining on the narrow ridge underneath LesSchuh. Note that the illustration solely illustrates the concept of a toeing-out strategy, and does not display the actual angles or scaling.

#### VI The center of pressure butterfly pattern with different shoes

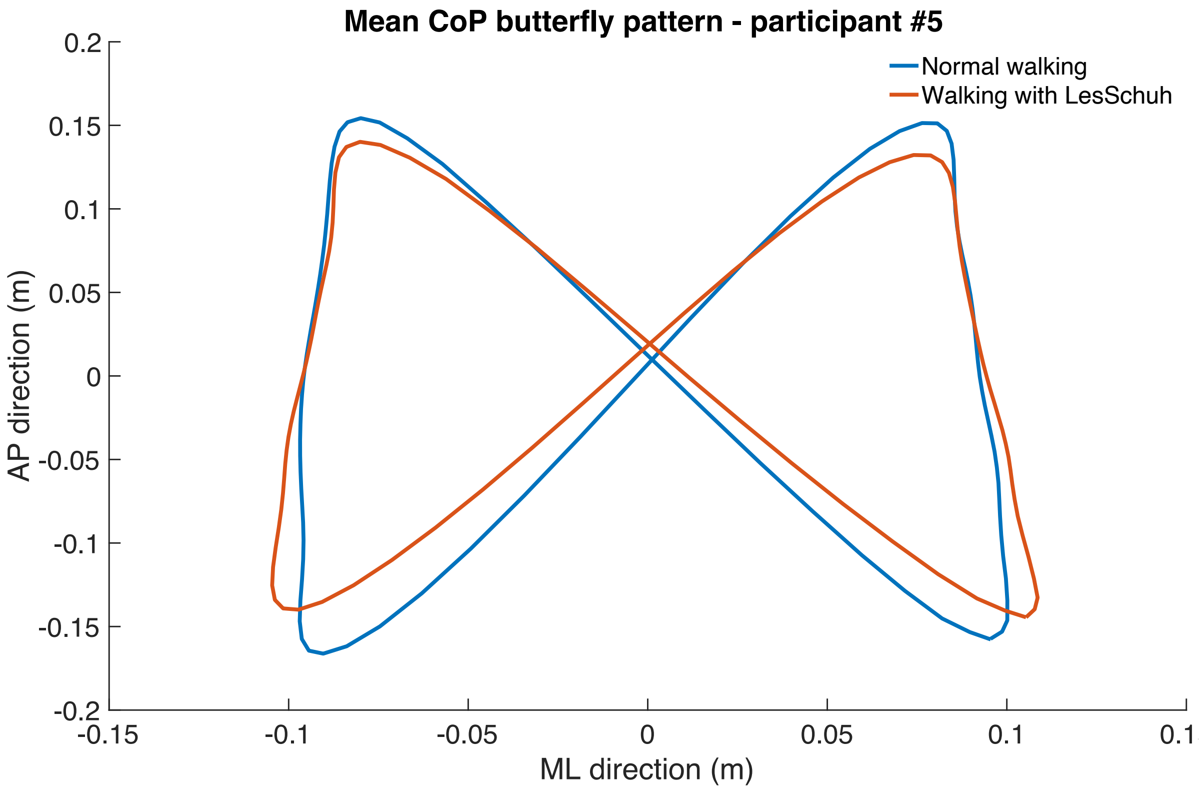


SVI.Fig1. Center of pressure butterfly pattern. During treadmill walking the center of pressure pattern shapes like a butterfly. This pattern can be used for gait event detection (Roerdink et al., 2008). For this figure, the mediolateral (ML) and anteroposterior (AP) coordinates were time-normalized from heelstrike-to-heelstrike, and then averaged to get the mean CoP trajectory in the force plate's coordinate system. Before plotting we subtracted the mean from each CoP trajectory. The resultant plot for participant 5 as an example, illustrates the similarity in butterfly-patterned-shape for normal walking (blue) and walking with LesSchuh (red).
